# baRcodeR with PyTrackDat: Open-source labelling and tracking of biological samples for repeatable science

**DOI:** 10.1101/457051

**Authors:** Yihan Wu, David R. Lougheed, Stephen C. Lougheed, Kristy Moniz, Virginia K. Walker, Robert I. Colautti

## Abstract

Repeatable experiments with accurate data collection and reproducible analyses are fundamental to the scientific method but may be difficult to achieve in practice. Several flexible, open-source tools developed for the R and Python coding environments aid the reproducibility of data wrangling and analysis in scientific research. In contrast, analogous tools are generally lacking for earlier stages, such as systematic labelling and processing of field samples with hierarchical structure (e.g. time points of individuals from multiple lines or populations) or curating heterogenous data collected by different researchers over several years. Such tools are critical for modern research given trends toward globally distributed collaborators using higher-throughput technologies. As a step toward improving repeatability of methods for the collection of biological samples, and curation of biological data, we introduce the R package *baRcodeR* and the *PyTrackDat* pipeline in Python. The *baRcodeR* package provides tools for generating biologically informative, hierarchical labels with digitally encoded 2D barcodes that can be printed and scanned using low-cost commercial hardware. The *PyTrackDat* pipeline integrates with *baRcodeR* output to build a web interface for sample management and tracking along with data collection and curation. We briefly describe the application of principles from *baRcodeR* and *PyTrackDat* in three large research projects, which demonstrate their value to (i) help document sampling methods, (ii) facilitate collaboration and (iii) reduce opportunities for human errors and omissions that could otherwise propagate through downstream data analysis to compromise biological inference.

## Introduction

The increasing use of high-throughput methods for data collection and popularity of large collaborative research projects in biology poses challenges for researchers tracking samples and their associated data across field and laboratory experiments. New standards and tools for documenting data analysis have developed alongside rapid advancements in computation and global communication, aimed at improving repeatability, reproducibility, accuracy and accountability of published scientific research. These methods are likely to become more prevalent as major funding bodies move toward principles and policies stressing well-developed data management plans (DMPs), as demonstrated in the Draft Tri-Agency Research Data Management Policy (Government of Canada), Article 29 of the ERC Multi-Model Grant Agreement (European Commission), and the NSF open data policy (National Science Foundation).

A push towards more transparent, reproducible data workflows for scraping, merging, cleaning, editing, error checking, pre-processing and curating data for subsequent visualization and statistical analysis (i.e. data wrangling) has inspired the rapid development of new tools. For example, R markdown and R notebooks with R Studio (RStudio Team, 2016) are now common tools to facilitate careful documentation of data visualization and statistical analysis in R (R Core Team, 2018). Jupyter Notebooks and Jupyter Lab (Kluyver et al., 2016) serve similar functions for reproducible analysis in Python (Oliphant, 2007). Such tools allow for fully reproducible data wrangling and analysis from a starting dataset, which was difficult or impossible with previous generations of proprietary spreadsheet databases and point-and-click statistical packages. In contrast, tools to improve transparency, accuracy, repeatability, and reproducibility at the earlier stages of sampling, data collection and curation have received relatively little attention, even though modern downstream management packages assume accurate sample tracking and recording of raw data and sampling details.

In many of the biological sciences, samples are collected under arduous field conditions that pose challenges for labelling, organizing, measuring and analyzing samples. A common approach is to quickly collect and preserve samples for later analysis. This often requires transport of samples to different locations, to different storage media or vessels, and/or subsampling for multiple analyses. An individual biological sample is often tied to a variety of heterogenous data such as locations, images, morphological measurements, behavioral assays, chemical analysis, as well as details about the sampling method itself, such as the identities of individuals collecting, processing, and measuring the samples. Heterogeneous data may be collected by different individuals or labs and stored in a variety of formats but are inherently linked by details about their collection such as sampling method, date and location, collector identity, and other metadata. Misinterpreted labels, cut-and-paste errors, typographical and scribing errors, and other problems that arise when humans associate samples with their downstream measurements and metadata can potentially go undetected and propagate through statistical analysis to compromise biological inference.

Asset tracking using digital barcodes can significantly reduce opportunities for human error by automating and/or acting as additional error checks on human tasks that are error prone (Copp et al., 2014). Indeed, this ‘industrial approach’ has been successfully adopted to improve cataloguing of natural history collections that may contain thousands to millions of specimens (Blagoderov et al., 2012). Perhaps the strongest evidence for the value of digital barcodes is their ubiquitous use in inventory tracking and sales in virtually every type of commerce on earth, ranging from perishable foods with short turnaround times to inventories in long-term storage facilities. In addition to reducing human error, digital barcoding technology can simplify data collection when combined with data collection software and a dedicated barcode scanner or image recognition applications for smartphone, tablet, or personal computers.

Numerous options are available for asset tracking using barcodes, many with integrated databases (e.g. ManageEngine, Pulseway, Asset Panda, GoCodes, OpenLab Framework). However, most of these solutions are proprietary and can be expensive (but see List et al., 2014). These programs prioritize ease-of-use, employing a graphical-user interface (GUI) that are relatively easy to learn but can limit options for customization, automation and integration with downstream data wrangling and analysis. Whereas numerous statistical packages exist that improve transparent and reproducible analysis (e.g. in R and Python) compared to point-and-click statistical software, we know of no analogous options for sample labelling and data collection. Integration of such options within the R and Python coding environments would facilitate documentation of data management from initial collection through to final analysis for more repeatable and reproducible research.

Here we introduce an integrated programming solution using the *baRcodeR* package in R with the *PyTrackDat* pipeline for labelling and tracking biological samples as well as collecting and curating their associated data. Our goal was to establish a workflow that is (i) open-source, (ii) based in R and Python, (iii) flexible, (iv) inexpensive to implement at scale, and (v) developed specifically for integration within a transparent and robust data management plan for repeatable and reproducible science. To demonstrate the utility of the software and the implementation of the workflow, we include three current case-use studies at different stages of development.

## Methods & Results

Initial surveys of commercially available digital barcode systems suggested that they were expensive or impractical for our needs. For example, although grocery store hand-held scanners are cheap and robust enough for fieldwork, commercial inventory software could not be easily adapted to suit data collection in the field. Our solution was to write the R package *baRcodeR* (Wu and Colautti, 2018a) and to extend its functionality with *PyTrackDat* a **Py**thon data pipeline for sample **track**ing and **dat**a collect we call (https://github.com/ColauttiLab/PyTrackDat).

### 1. baRcodeR

The *baRcodeR* package for R has a detailed tutorial-style vignette (Wu and Colautti, 2018b) and a 2-page ‘cheat-sheet’ summary on FigShare (Wu and Colautti, 2018c). The latest release can be installed directly in R using the command:

~~~
**> install.packages(‘baRcodeR’)**
~~~

A pre-release of the latest features can be installed from GitHub (https://github.com/yihanwu/baRcodeR), either by manually downloading and installing the binaries or using the devtools package:

~~~
**> devtoools::install_github(‘baRcodeR’)**
~~~

Briefly, *baRcodeR* was developed as an open-source tool for creating digital 2D barcodes in R, making several design choices based on our experiences collecting and managing biological samples. Three main commands are available: (i) *uniqID_maker()* or (ii) *uniqID_hier_maker()* are functions to generate unique ID codes, followed by (iii) *create_PDF()* to output a portable document file (PDF) containing both text labels and 2D barcode images laid out in a customizable format suitable for consumer-grade printers. These functions can be run directly from the command line or via the *baRcodeR* graphical user interface (GUI) available from the ‘Addins’ menu in R Studio. The command-line version also includes a more user-friendly interactive mode implemented with the parameter *user=T*.

#### Unique identifier codes

To generate standardized and biologically informative ID codes for biological samples, *uniqID_maker()* or *uniqID_hier_maker()* are used for sequential or hierarchical text labels, respectively. The *uniqID_maker()* command is particularly useful to quickly produce sequential labels with the same user-defined prefix string (e.g. *Ex-1*, *Ex-2*, *Ex-3,*…). Additionally, it is possible to pass any sequence of numbers into *uniqID_maker()*. For example, the user can specify alternating numbers (e.g. *Ex-001*, *Ex-003*, *Ex-005,*…):

~~~
**> uniqID_maker(string = ‘Ex’, level = seq(1, 6, 2))**
~~~

or other custom sequences (e.g. *Ex-010*, *Ex-014*, *Ex-018*, *Ex-020*, *Ex-040*):

~~~
**> uniqID_maker(string = ‘Ex’, level = c(seq(10, 20, 4), 20, 40))**
~~~

or randomly ordered sequences:

~~~
**> uniqID_maker(string = ‘Ex’, level = sample(1:10, replace = F))**
~~~

The *uniqID_hier_maker()* command expands on *uniqID_maker()*, allowing for the creation of hierarchical labels with different user-defined strings and numeric sequences for each level of the hierarchy. This is useful, for example, to generate labels for replicated populations, genetic lines nested within populations, and repeated sampling or subsampling of individuals at different time points, with randomly assigned treatment categories. For example, one can quickly define labels for ten replicate individuals, each from one of three lines, with subsamples of two different tissue types with the command:

~~~
**> uniqID_hier_maker(hierarchy = list( c(‘line’, 1, 3),**
~~~

~~~
      **c(‘ind’, 1, 10), c(‘sbsmpl’, 1, 2)))**
~~~

The examples of *uniqID_maker()* and *uniqID_hier_maker()* use non-interactive mode by default, but can also be run interactively in the command line with the *user = T* parameter. In this mode, the user is prompted for input:

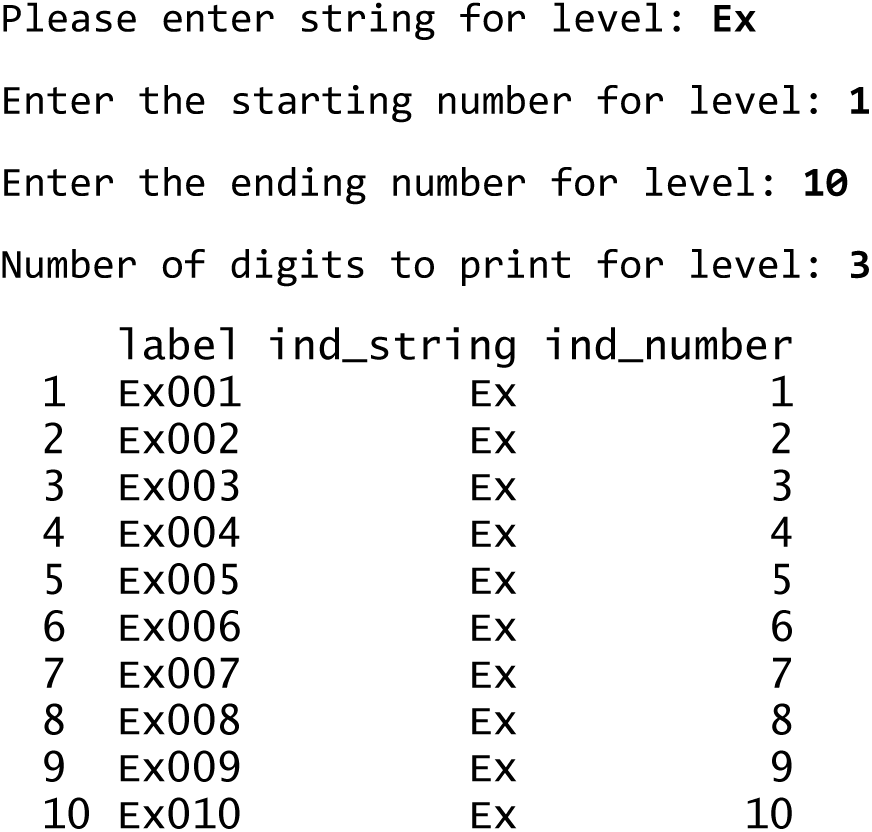

Regardless of mode (i.e. interactive or non-interactive), the output of *uniqID_maker()* and *uniqID_hier_maker()* is a *data.frame* object containing (i) a vector of unique IDs and (ii) additional columns corresponding to each level of the sampling hierarchy. Like any *data.frame* object in R, the output can be renamed with *names()* function. The output object can also be saved to a standard text file using the *write.table()* or *write.csv()* functions. These text files can then be imported into a spreadsheet or database software, or integrated with *PyTrackDat* (see section 3, below) as part of a documented research workflow.

#### Printable barcodes

A *data*.*frame* containing a vector of ID labels is required input for the *create_PDF()* function. The input *data.frame* may be created by one of the functions above or supplied by the user, for example by entering ID codes manually into a spreadsheet, exporting to CSV, and then importing the CSV file into R using the *read.csv()* command.

The output of *create_PDF()* is a printable PDF file containing human-readable (i.e. plain text) IDs and 2-D digital barcode images. These labels can be printed on consumer-grade printers and scanned with standard barcode scanners or image recognition software for cellphones or other devices(Fig. 1). The specific print layout can be customized using spacing parameters in *create_PDF()*, but default parameters create a PDF output file that will fit S-19297 labels (ULINE.ca or ULINE.com). This layout was chosen because the S-19297 labels are weatherproof vinyl labels (80 per page) that can be printed on a standard laser printer. Our own tests demonstrate that printed labels do not fade or degrade even after two years of storage at -80°C or three years of exposure to sunlight in outdoor field experiments. Standard laser printer toner is similarly robust to ultraviolet light and freezing to ultralow temperatures. The label glue is less robust but transparent packing tape can be used to better secure the labels without affecting their function.

**Figure 1.**
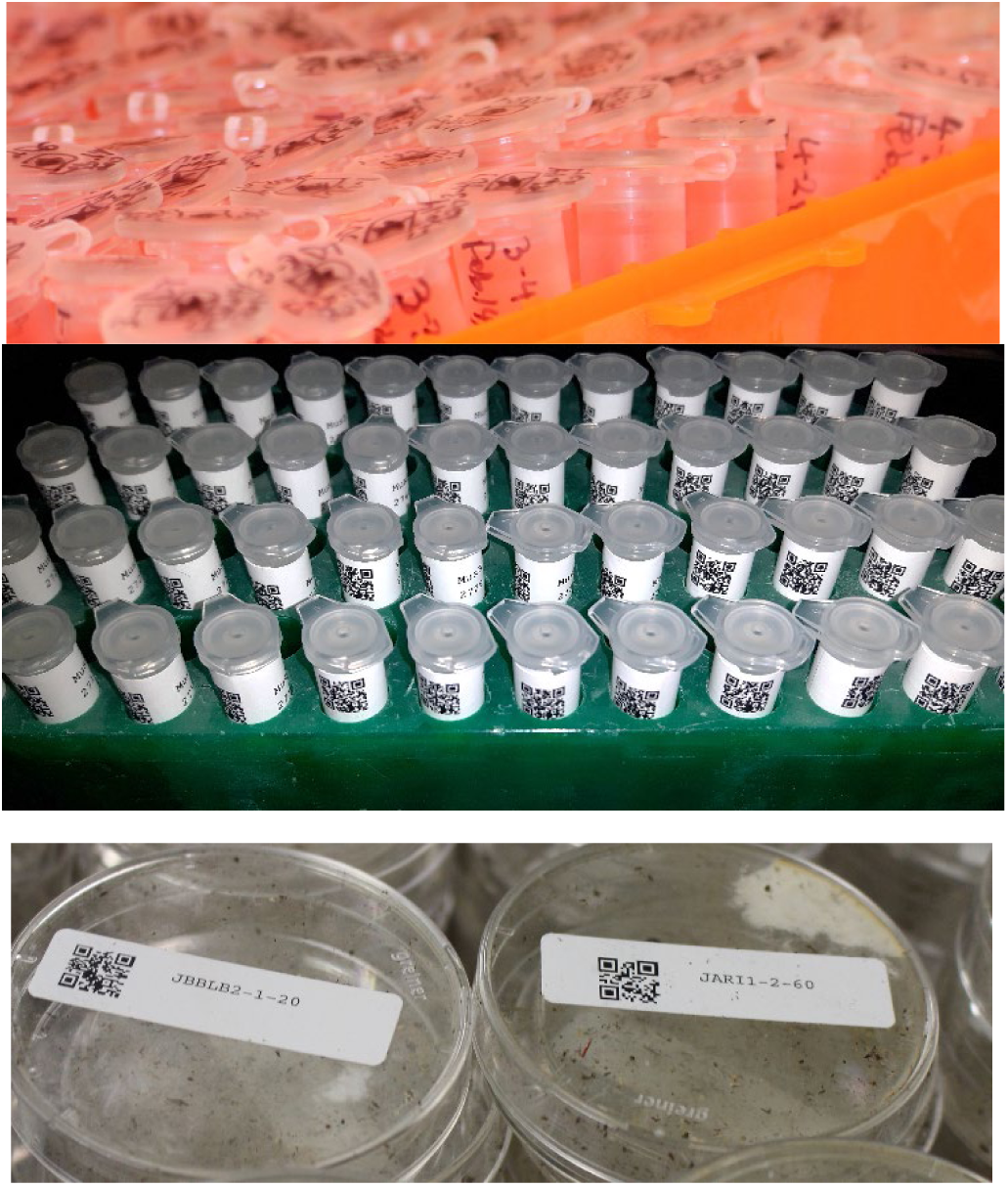
Examples of conventional, hand-labelled tubes (top panel), compared with tubes (mid panel) and petri dishes (lower panel) labelled with digital barcodes and human-readable text labels created using the *baRcodeR* package.

The *create_PDF()* function can be run interactively with the *user = T* parameter or in default non-interactive mode with user-defined parameters for custom label layouts and other parameters like file name and font size. One parameter of particular note is the error correction level (L, M, Q or H), which represent a trade-off between barcode size and redundancy, ranging from larger, high-redundancy (H) to smaller low-redundancy (L) 2-D barcodes. H-codes can still be recognized by scanners after losing up to 30% of their surface while L-codes labels are less robust but can be printed at a small size. Custom page layout, label layout and printing options can be specified in by passing in arguments through *create_PDF()* to *custom_create_PDF().* See the *baRcodeR* vignette for details (Wu & Colautti 2018b).

#### Graphical user interface

In addition to the text commands reviewed above, a graphical-user interface (GUI) is available after installing and loading the *baRcodeR* library (RStudio, 2015). Details of the GUI are provided as a quick-start guide in the README file on the CRAN repository (Wu & Colautti 2018a; direct link: https://cran.r-project.org/package=baRcodeR/readme/README.html). The GUI add-in is a *shiny* application (Chang et al., 2018) that can be run directly from the *baRcodeR* GUI under the ‘Addins’ menu on the RStudio toolbar. Three tabs in the GUI window correspond to three main functions: ‘Simple Label Creation’ calls *uniqID_maker()*, ‘Hierarchical Label Creation’ calls *uniqID_hier_maker()*, and ‘Barcode Generation’ calls *create_PDF()*. Labels generated from the first two tabs are previewed before final generation and will be automatically saved in the R working directory upon creation. All advanced parameters for PDF layout are similarly available under the “Barcode Generation” tab. Users can also preview the layout of individual labels before creation. Code snippets are shown for user-specified parameters and can be exported and archived as part of a repeatable experimental design and reproducible analysis.

In addition to using *create_PDF()* to generate printable barcodes, users can load saved labels into other barcode printers, such as those that print directly on sample tubes, plant tags, and other material. For example, the TubeWriter 360 (https://tubewriter.com/) and TubeMarker 2 (https://www.4ti.co.uk/) can generate and print barcodes from standard text files, allowing users to create ID codes in *uniqID_maker()* or *uniqID_hier_maker()* and saving the output to a text file using the *write.table()* or *write.csv()* functions in base R.

Once a *data.frame* or CSV file is created, additional headings can be added for data entry. Manual data entry is inherently susceptible to human error, but enforcing and restricting entry formats can help to ensure repeatability of scientific inference. A variety of software is available to help reduce errors in data entry, ranging from fully-featured propriety software suites with optical recognition of handwritten data such as Viking Data Entry (http://vikingsoft.com), to open-access programs such as Epidata (Lauritsen and Bruus, 2003) and OpenRefine (http://www.openrefine.org). Microsoft Excel, Libreoffice Calc, and Google Sheets also have data form creators for restricting data entry types. Within RStudio, interactive data entry can be performed using the *editData* add-in (Moon 2017), which can be appended to the biological sample *data.frame* created from *baRcodeR*. Alternatively, users may wish to implement the semi-automated *PyTrackDat* pipeline to build an online relational database as described in the next two sections.

### 2. Data Standards

Perhaps the simplest implementation of a robust data collection workflow is a 2-dimensional spreadsheet (i.e. data table) containing *n* rows by *c* columns. This format has several important properties and considerations, as shown in the example in Figure 2, outlined in Table 1, and discussed below (see also Borer et al., 2009; British Ecological Society, 2014). In this format, each column is a characteristic or measurement and each row is (usually) an individual, with each individual cell containing a single value. To facilitate reproducible analysis, spreadsheet data should not contain empty spaces or formatted text (e.g. colour, bold, italic, underline). Rather, individual values can be repeated, and missing values can be encoded as needed (e.g. NA or NULL). Variable names should be chosen carefully to be as short as possible but still informative. A good strategy for more complicated names is to remove vowels or to use short forms separated by capital letters, underscore or period. For example, a column containing height measurements on day 5 of an experiment might be named HtD5 or Hght.d5. Quotation marks, spaces, commas, semicolons and other common separator characters should be avoided as these are often used by programmers and software to parse data into separate rows or columns. Where necessary, for example in a ‘notes’ column, these characters can be used in a phrase denoted by quotation marks (e.g. “This is a note; it can be included, if needed, in a ‘notes’ cell”).

**Figure 2.**
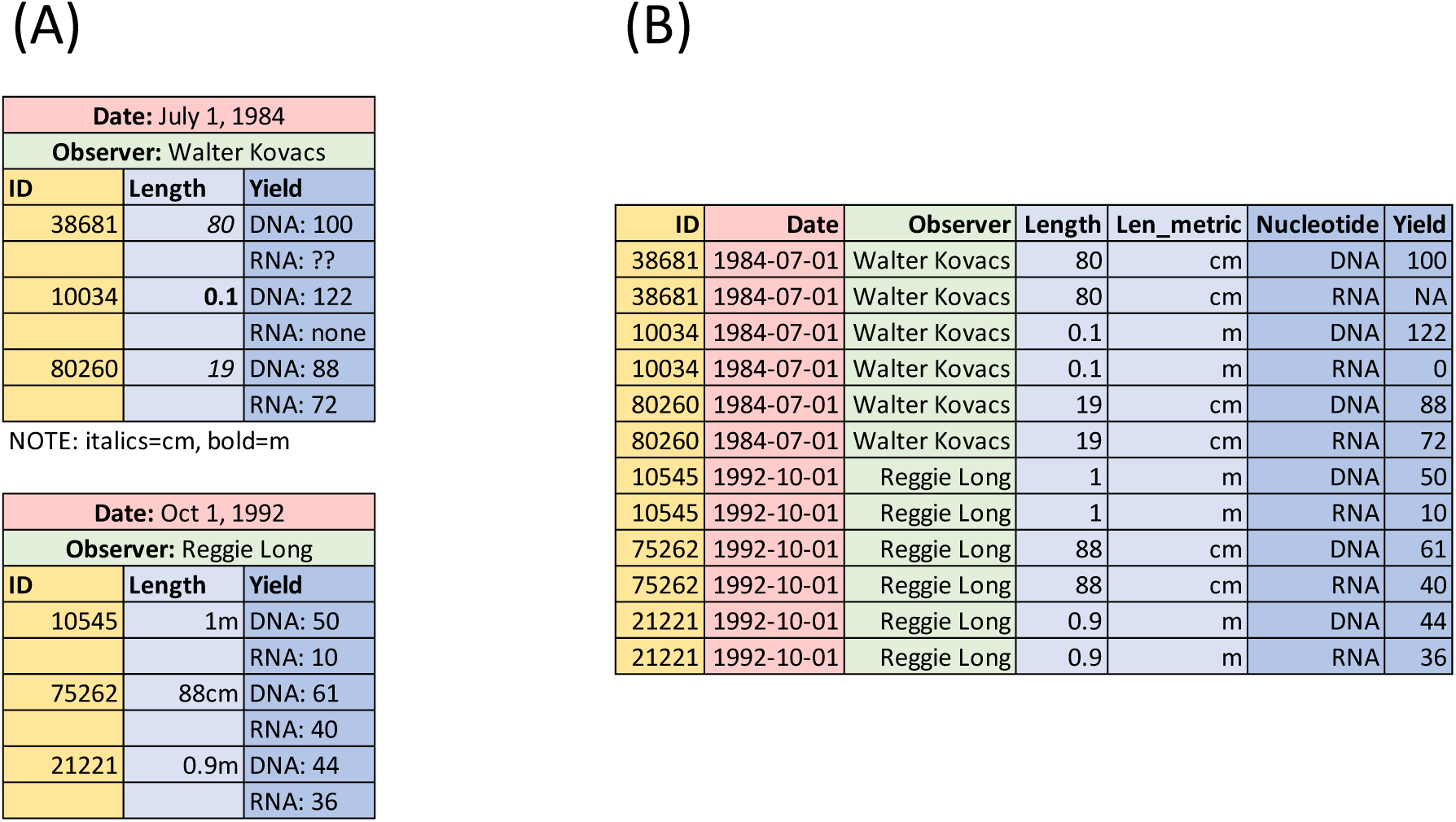
Example of (A) common errors in data management and (B) corresponding rearrangement of the same data to simplify reproducible data wrangling and analysis. Note that colours are added to show link between data in A and B and do not appear in the final text-based file (e.g. TXT, CSV, TSV).

**Table 1.** . Summary of recommended data standards

| Standard | Example |
| --- | --- |
| Consistent naming | Use a single value for representing a female individual, rather than inconsistent use of, for example, 'F', 'f', 'Female' and 'female'. |
| Appropriate data types | Integer (42), Float (3.14), String ('cat'), Boolean (True/False), or Date (YYYY-MM-DD) where appropriate. |
| Homogenous data types | Don't mix data types in a single column. For example, if you measured in cm and mm, use one integer column for the measurement and a separate string column for the unit of measurement. |
| Human readable, non-proprietary formats | Data file can be opened using a standard text editor. For clarity, these files are typically saved with the suffix *.txt*, *.csv* (comma-separated) or *.tsv* (tab-separated). |
| Appropriate null coding (no empty cells) | Use 0 for observed zeros but NA<sup>1</sup>, NaN, or NULL for missing data. |
| Appropriate variable names | Short but informative variable names, perhaps combining multiple words (e.g. flowering time) using short-forms with a period (flwr.t), underscore (flwr_t) or capital letter (flwrT). |

<sup>1</sup>NA is the preferred value for missing data in R.

In addition to eliminating empty spaces and conforming to the 1 cell = 1 characteristic rule, each column should be homogenous; that is, a column should contain only a single type of data. Data types are typically *integers* (whole numbers between -4324 and 10000002), *floating point numbers* (non-integer numbers), *boolean values* (just two states: True or False; 1 or 0) or *strings* (alphanumeric characters). As shown in Figure 2, each column should be homogeneous – that is, all entries contain the same type of data. For example, if length measurements were made in *cm* and *m*, then the measurements ‘1m’ and ‘10cm’ should be separated into integer and string columns to avoid heterogenous data, as shown in Figure 2. Using different values for the same entity also creates undesirable heterogeneity in a data column, for example alternating between ‘F’, ‘f’, ‘Female’ and ‘female’, which are interpreted as different values by analysis software. Notes about individual observations (i.e. rows) or the overall dataset (i.e. metadata) should not be hard-coded as stand-alone rows in datasheets, as shown in Figure 2a, but instead encoded as columns or included in a separate file called readme.txt, notes.txt, metadata.txt, or something analogous. A metadata file (e.g. metadata.txt) should include a list of each column and all variable names, with brief descriptions and units of measurement. Similarly, associated files such as images or DNA sequences (e.g. FASTA or FASTQ) can be indexed with reference columns containing file names or ID codes that match to corresponding image file names or DNA sequence IDs. Finally, data should be saved in a non-proprietary, human readable text file format to ensure longevity (e.g. TXT, TAB or CSV file).

In addition to the basic standards outlined above, multiple schema are available to standardize and share data. For example, Ecological Metadata Language (Fegraus et al. 2005) is a data schema which puts no restraints on the kinds of data collected but requires comprehensive documentation of data through metadata standards. Darwin Core (Wieczorek et al., 2012) is another schema that uses each row as a record and has a variety of standardized column names. The resulting data package can then be uploaded and shared on online repositories such as DataONE, Dryad, and the Knowledge Network for Biocomplexity (Andelman et al., 2004; Greenberg, 2009; Michener et al., 2012). Biodiversity repositories such as the Global Biodiversity Information Facility accept Darwin Core datasets with metadata files.

### 3. PyTrackDat Pipeline

The *PyTrackDat* pipeline (https://github.com/ColauttiLab/PyTrackDat) contains two main Python scripts: The first script analyzes text-based data files, such as the *data.frame* object generated from *baRcodeR*, to generate a design file as described below. The second script uses a design file to generate a web application for sample tracking and data collection. Although designed to integrate with *baRcodeR*, both scripts in *PyTrackDat* can be run independently on user-defined input files.

The first part of the *PyTrackDat* pipeline is an optional *DataAnalyzer* script that assists with implementation of the data standards outline in the previous section. It can be run with a single CSV-formatted input file, or a series of CSV files linked by one or more column names (e.g. sampleID). In the latter case, the code will remap fields (i.e. columns) from multiple data tables into a single, more detailed database format with associated metadata. This format specifies field properties such as data type, acceptable values, and whether fields can contain NULL values (Fig. 3). The design file should be inspected manually using a text editor to look for errors (e.g. inconsistent data types) and to add descriptions about the data. The edited design file serves two main purposes: (i) archiving data types and other metadata that isn’t easily stored in a spreadsheet format; (ii) rigidly defining fields and links among data tables to create a relational database.

**Figure 3.**
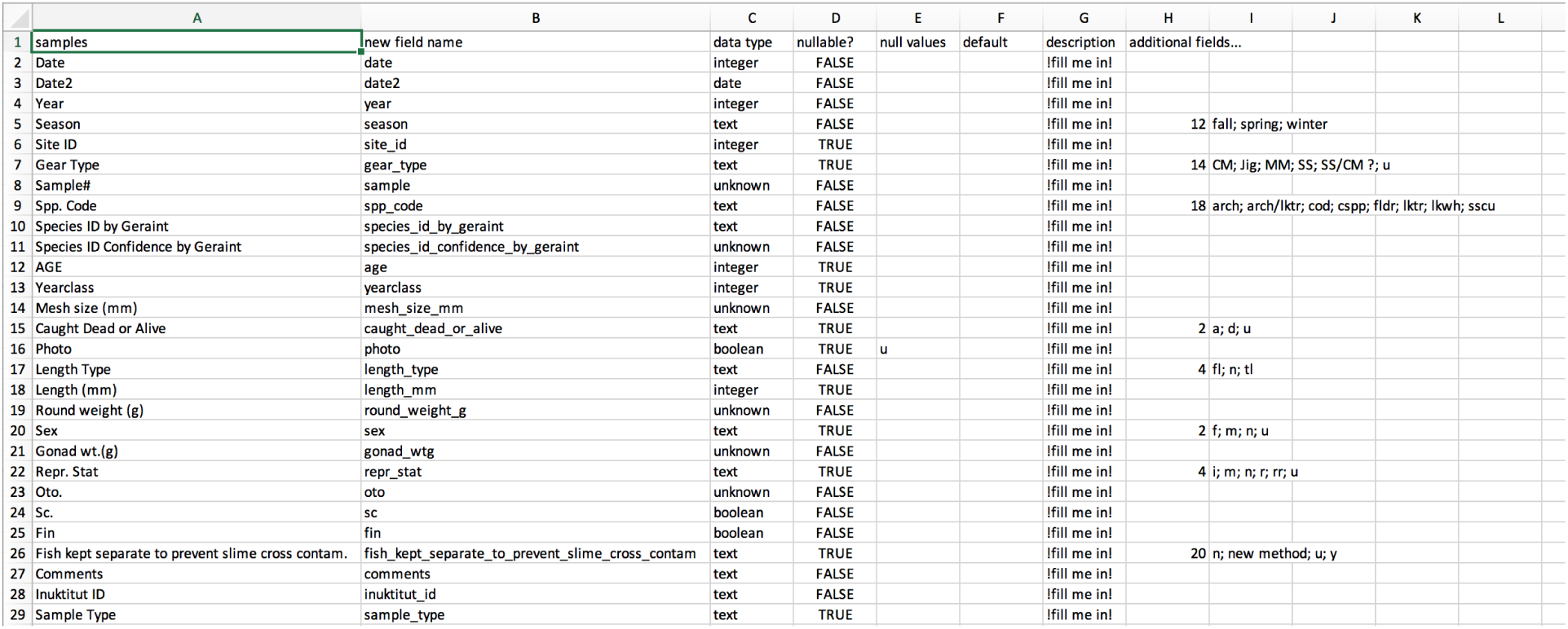
A sample design file generated from the *DataAnalyzer* script in the *PyTrackDat* pipeline. Data types are automatically inferred for each data column in the CSV file, with ‘unknown’ indicating fields that should be investigated by the user. The number of different options for text variables are shown under ‘additional fields’, along with some examples.

The second script is *WebBuilder*, which generates a web-app for data collection based on input from a design file. The design file can be a completed version of the file generated from the *DataAnalyzer* script or a de-novo design file written by the user using a text editor. Based on this input, the second script in the *PyTrackDat* pipeline creates a zipped folder containing the packaged web application, ready to be deployed (see README.md at https://github.com/ColauttiLab/PyTrackDat).

### 4. Implementation

To limit errors while improving repeatability of methods and facilitating reproducibility of downstream analysis, we suggest two alternative strategies for data collection and curation. For small projects with one or a few investigators working together, such as a student thesis project or for a focused technical report, a basic data management framework proposed above may be appropriate. Further considerations are needed for large, complex databases used by multiple investigators from different research groups. Although it is theoretically possible to encode everything in a single spreadsheet format, this may require frequent edits by multiple users, resulting in an unstable dataset. Instead, data can be organized into separate files linked by identifying columns (e.g. sample, collector, location, date). These are referred to as ‘relational databases’ and many software packages are available to handle such data, including dBase (dBase LLC, Binghampton; http://www.dbase.com), FileMaker Pro (FileMaker, Santa Clara; http://www.filemaker.com), Google Fusion Tables (Alphabet, Santa Clara, http://drive.google.com), LibreOffice Base (The Document Foundation, Berlin, http://www.libreoffice.org), and MySQL (Oracle, Redwood City, http://www.mysql.com).

To facilitate creation and collection of both ‘basic’ and ‘relational’ data, *baRcodeR* and *PyTrackDat* were designed to facilitate sample management and data collection according to the standards outlined in Section 2, above. These standards include generation of code and text files to document sample ID, data management decisions, and data characteristics as an important but underappreciated step in open, repeatable, and reproducible science. The utility of our design philosophy can best be illustrated by three case studies, outlined briefly below, which involve collaborative projects spanning large geographical regions and involving dozens to hundreds of collaborators studying microbes, plants and animals using a large variety of conventional and cutting-edge high-throughput technologies. These projects are illustrative as they represent three different stages of development: Example 1 predates *baRcodeR* and inspired many of its features, example 2 was co-developed *baRcodeR* and *PyTrackDat*, and example 3 is a new project informed by these standards and inspiring further refinement.

### Example 1: GGMFS

The ‘Global garlic mustard field survey’ (GGMFS) is a coordinated, distributed experiment in evolutionary ecology involving >150 collaborators from 16 countries spanning Europe and North America (www.garlicmustard.org). The overall goal of the project is to focus research effort on a single focal species, *Alliaria petiolata* (garlic mustard), to better understand how natural and anthropogenic processes affect genotype, environment and genotype-by-environment effects on phenotypic traits and ecological interactions at local (< 10 m) to global spatial scales. Details of the project, including protocols and design philosophy are discussed elsewhere (Colautti et al., 2014). Here we elaborate on the logistical challenges that arose from this large collaborative project and how it informed the design philosophy applied to *baRcodeR* and *PyTrackDat*.

The project began with a relatively simple field sampling protocol to collect seeds and to measure plant size and fecundity, population density, herbivory and pathogen damage, as well as characteristics of the surrounding environment (protocol https://doi.org/10.6084/m9.figshare.729274). To link seed collections with their associated data, we implemented a sample naming standard using the general format:

~~~
**2010 JDNYC1-3-20**
~~~

Where 2010 is the year of collection, JD are the initials of the collector, NYC is a 3-letter location code provided by the collector, 1 is the first population sampled at that location, and 3-20 denotes a sample from 3m and 20cm along a 10m transect. Participants also took canopy photos at 3 points along the transect, and the resulting image files were saved with file names incorporating the population code and transect location. Although a standard protocol was implemented, minor differences among the >150 different academic participants posed challenges to collect and curate standardized data. To facilitate data entry, we used an online survey service (surveymonkey.org). A web-based portal simplified data aggregation but in several cases the participants deviated from established protocol in their choice of measurement format. This led to a variety of ambiguous entries that took three years to correct.

Perhaps the most problematic issue was the entry of GPS coordinates, which were entered in every format available: DMS: 41°24′12.2″N 2°10″26.5″E, DMM: 41 24.2028, 2 10.4418, DD: 41.40338, 2.17403, and several confusing hybrids (e.g. 41.24.2028). To correct these issues, we wrote custom code in R using regular expressions and sent hundreds of emails to clarify contributed data. Enforcing homogeneous data standards from the beginning would have saved hundreds of hours of work.

Current studies associated with the project use the archived seed collections for laboratory and field experiments measuring a variety of phenotypic traits, genotyping using high-throughput sequencing methods, and soil microbial feedbacks. The use of a standard ID code allows a quick link back to any of the original field measurements. Once published, data from current and future studies can be linked via the biologically-informed ID format and quickly accessed for further analysis using the relational database framework described above.

### Example 2: TSFN

From the inception of the Genome Canada project, “Towards a sustainable fishery for Nunavummiut” (TSFN) it was understood that this collaborative effort between more than half a dozen research labs would be necessary to successfully track thousands of samples collected over an area of 2500 km^2^ under the challenging conditions of the high Arctic in all seasons. Selected data must be accurate, repeatable and reliable for upload to a digital geographic Atlas, providing a rich and accessible resource for other researchers and Inuit community members. We posited that digital barcoding would provide the only reliable means to track each and every fish sample (fin clips, right and left otoliths, organ biopsies, gonads, skin mucous, intestinal sections, parasites, scales, muscle for contaminant assays and genetics), as well as the metadata associated with each sample (e.g. geographic location, date, name of fisher, net size, net set hours, species, sex, weight, length, photograph). Efforts to purchase software were abandoned as we could not find an affordable solution that was flexible enough to quickly generate biologically meaningful sample ID tags and to accommodate a diverse range of label sizes for various sub-sampling containers.

Using *baRcodeR*, we created fish-specific ID codes with 2D barcodes printed on waterproof paper and vinyl labels in convenient sizes such as 10 × 5 cm and 4.5 × 1.3 cm; Uline, Milton, ON). These were used to label cryotubes, centrifuge tubes, and a range of sizes of re-sealable plastic bags and manila envelopes (Fig. 4). These sample containers were aggregated into a large plastic bag (27 x 27 cm), affixed with the fish-specific barcode mounted on a white sheet bearing a 9.5 cm-scale line beneath the lower barcodes. The container with scale bar was included in photographs of the fish as a redundant check on the accuracy of the reported length measurements. Barcodes were initially hand-scanned (reader CR1421-PKU, Code Corp.) directly into a spreadsheet in Microsoft Excel (ver. 16.16.1) with descriptors added manually. However, data heterogeneity resulting from project participants who were not familiar with the recommended data guidelines (Table 1) prompted development of *PyTrackDat*. For example, although the length of each fish is routinely measured from the head to the center of the tail fork (fork length), some lengths were reported as head to the end of the squeezed tail (total length); this necessitated the addition of a separate column for total or fork length designations to avoid heterogenous data (see Section 3 guidelines on data standards). The use of the *PyTrackDat* pipeline to create an online data collection and tracking system facilitated data entry according to above standards by multiple users. Data are currently curated by a small number of experienced users before sharing with other collaborators.

**Figure 4.**
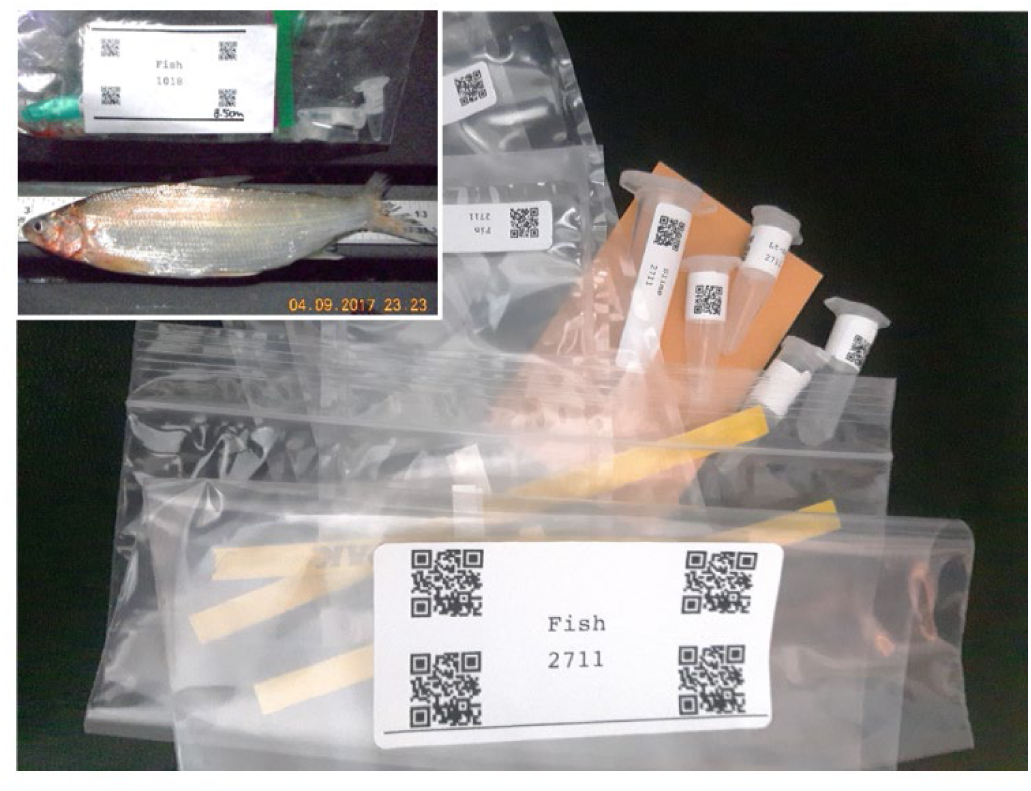
Sampling kit demonstrating the use of *baRcodeR* to improve data integrity and repeatability of sampling methods. A sample bag for each fish is shipped to the field; each bag contains labelled containers for various tissue subsamples. In this example, separate labels were created with *barRcodeR* to encode the same fish ID #2711 into the bag containing the fish, as well as all subsamples associated with it. Inset shows a cropped archival picture of fish #1018, which visually links a fish ID code to its collection date and time, fish length and species ID.

### Example 3: BEARWATCH

The BEARWATCH project seeks to develop a community-driven program to facilitate ongoing monitoring of polar bears across the Canadian Arctic, combining Inuit traditional knowledge with a toolkit for characterizing key aspects of population health. In addition to extensive consultation with northern communities and collation of existing traditional knowledge, Phase 1 of the project involves creating a baseline map of genetic structure by genotyping thousands of individuals harvested by indigenous hunters or biopsied by territorial governments over several decades. Subsamples from archived tissue arrived either fixed with 95% ethanol in Eppendorf vials, dried in an envelope (skeletal muscle), or as a preserved biopsy plug (with associated fat, skin, hair, and muscle). All have either a NT (Northwest Territories) or NU (Nunavut) field sample number previously assigned by territorial governments (e.g. NT_100177, NT_100178, NU_L26303, NU_L35160). Associated metadata include geographical coordinates (decimal degree latitude and longitude), management unit (one of 19 worldwide indicated by a 2-letter code; e.g. Gulf of Boothia=GB, Lancaster Sound=LS; see Obbard et al., 2010), collection date (Year-Month-Day), sex (M, F), and age (Adult, Subadult, 2Yearling). Upon receiving samples to our Biosafety Level 2 laboratory, each is assigned a barcode generated using *uniqID_maker()* (Um-000001, Um-000002, Um-000003). The prefix ‘Um’ is for *Ursus maritimus*, followed by 6 digits anticipating a continuous monitoring system that can accommodate continued influx of samples for the foreseeable future.

We are using a reduced-representation genotype-by-sequencing approach (double-digest RAD-Seq; Peterson et al., 2012) to generate large panels of Single Nucleotide Polymorphism (SNP) data that will be used for quantifying population structure and forensic genotyping. From Illumina sequencing runs, each individual bear will have an associated FASTQ file containing millions of individual sequence short reads, with details on instrument, run date, read quality, etc. As these data are too extensive to store in our master database, unique FASTQ file names (barcode_FASTQ.txt) will be listed as a string within one variable field for each bear referencing the pertinent FASTQ files stored elsewhere on our servers.

Phase 2 of the project will analyze contaminant loads, stable isotope profiles, and diet using DNA barcoding of polar bear scat collected across the Canadian Arctic (an area larger than Western Europe) by Inuit hunters and other agencies. Polar bear DNA from each scat sample will also be analyzed using a genetic toolkit we are developing to determine the sex and genetic identity from a subset of 182 bi-allelic SNPs identified in Phase 1. Each scat sample will again have associated metadata that includes geographical coordinates, management unit, and collection date. Each scat will be subsampled four times for separate genomic, contaminant, stable isotope, and contaminants analyses to be done in laboratories at three different institutions. We will use *uniqID_hier_maker()* to generate unique identifiers for tracking individual subsamples based on a pre-defined numbering system (1-genomics, 2- contaminants, 3-stable isotopes, 4-diet) – e.g. Um-000113-1, Um-000113-2, Um-000113-3, Um-000113- 4).

As this project ultimately envisions continued community-based scat sampling over decades, we anticipate receiving multiple scat samples belonging to the same individuals obtained in distinct seasons, years, and locations – the range of individual polar bears can encompass areas > 100,000 km^2^. (Ferguson et al., 1999). This will allow us to assess movement patterns and changes in contaminant load, diet, and trophic level for individual bears over time and space. However, we will not know initially whether we have sampled the same individual or close relatives until we have created the SNP profile, and either tested for genotype matches (e.g., Jin et al., 2017) or estimated relatedness using a program like SNPRelate Version 1.14.0 (Zheng et al., 2012). A separate data table will contain a pairwise-relatedness matrix of all scat samples, which will allow for later grouping of barcodes linking individuals based on a threshold similarity in the relatedness matrix, as well as samples representing likely relatives (e.g. full-sibs, half-sibs, parent-offspring). Overall, this project demonstrates the utility of *baRcodeR* with *PyTrackDat* when designing sampling strategies that must link heterogeneous data across researchers and collection dates, with the flexibility to add new data fields based on analyses that occur long after samples have been collected and processed.

## Conclusion

The rigour of scientific theories in the biological sciences rest on fundamental concepts of repeatable experimental methods coupled with reproducible data analysis. Many key and fundamental results of published research in the sciences and social sciences are not reproducible (Baker, 2016), due at least partly to insufficient published detail about experimental methods, data collection and statistical analysis (Ioannidis, 2005). The use of open-source tools for documenting data wrangling and analysis in scientific studies are important steps towards improving reproducibility and transparency of data published research. To complement these tools, we have introduced *baRcodeR* and *PyTrackDat* to improve transparency and repeatability upstream of data wrangling and analysis. This includes documentation of sampling design, data collection, and dataset curation. Using examples, we have demonstrated how these tools are particularly useful to large-scale collaborative research projects in biology, studies using high-throughput methods for data collection, and more generally for researchers striving to improve the documentation, repeatability and reproducibility of their scientific endeavours.

## Acknowledgements

We thank E. Bao for contributions to baRcodeR code, Dr. P. V.C. de Groot for his efforts in the TSFN fish and BEARWATCH sampling as well as E. Jensen, R. Clemente-Carvalho, K. Flock, the Gjoa Haven, NU community residents, the Hunters and Trappers Association (HTA), the Government of Nunavut, Natural Sciences and Engineering Research Council (Canada) and large-scale Genome-Canada Projects, and these projects’ associated supporters. We would like to thank all sampling personnel that provided invaluable help in the field, along with our Inuit guides who were integral to the Arctic sampling efforts. We also thank. We further acknowledge that Queen’s University is situated on traditional Anishinaabe and Haudenosaunee territory and we are grateful to be able to be live and learn on these lands.

